# Localised expression of *OsIAA29* suggests a key role for auxin in regulating development of the dorsal aleurone of early rice grains

**DOI:** 10.1101/2021.03.04.434009

**Authors:** Mafroz A. Basunia, Heather M. Nonhebel, David Backhouse, Mary McMillan

## Abstract

Endosperm of rice and other cereals accumulates high concentrations of the predominant *in planta* auxin, indole-3-acetic acid (IAA) during early grain development. However, IAA signalling and function during endosperm development are poorly understood. Here, we report that *OsYUC12* (an auxin biosynthesis gene) and *OsIAA29* (encoding a non-canonical AUX/IAA) are both expressed exclusively in grains, reaching a maximum 5 to 6 days after pollination. *OsYUC12* expression is localized in the aleurone, sub-aleurone and embryo, whereas *OsIAA29* expression is restricted to a narrow strip in the dorsal aleurone, directly under the vascular bundle. Although rice has been reported to lack endosperm transfer cells (ETCs), this region of the aleurone is enriched with sugar transporters and is likely to play a key role in apoplastic nutrient transfer, analogous to ETCs in other cereals. *OsIAA29* has orthologues only in grass species; expression of which is also specific to early grain development. *OsYUC12* and *OsIAA29* are temporally co-expressed with two genes *(AL1* and *OsPR602)* previously linked to the development of dorsal aleurone or ETCs. Also up regulated at the same time are a cluster of MYB-related genes (designated *OsMRPLs)* homologous to *ZmMRP-1,* which regulates maize ETC development. Wheat homologues of *ZmMRP-1* are also expressed in ETCs. Although previous work has suggested that other cereals do not have orthologues of ZmMRP-1, our work suggests OsIAA29 and OsMRPLs and their homologues in other grasses are part of an auxin-regulated, conserved signalling network involved in the differentiation of cells with ETC-like function in developing cereal grains.

**Main Conclusion:** Non-canonical AUX/IAA protein, OsIAA29, and ZmMPR-1 homologues, OsMRPLs, are part of an auxin-related signalling cascade operating in the dorsal aleurone during early rice grain development.

## Introduction

The yield and quality of cereal grains is dependent on the coordinated regulation of endosperm development, uptake of photosynthate and production of storage molecules, and is highly susceptible to adverse environmental influences (Yu et al. 2015). In addition, the molecular and physiological events taking place during early endosperm development determine to a large extent the final grain size and weight in rice and other cereals (Mizutani et al. 2010; Fahy et al. 2018). There is a large body of literature reporting the involvement of plant hormones in the regulation of early endosperm development as well as its response to environment (reviewed by Basunia and Nonhebel 2019). However, little is known of the detailed role of hormonal signalling and how this influences key processes of endosperm cellularisation, differentiation of cells responsible for nutrient uptake or the expression of starch and storage protein synthesis genes.

Previous work in our laboratory has shown that a large increase in the auxin, indole-3-acetic acid (IAA), occurs during endosperm cellularisation, aleurone development and the initiation of starch production in rice grains, driven by strong up-regulation of key auxin biosynthesis genes, *OsTAR1, OsYUC9, OsYUC11* and *OsYUC12* (Abu-Zaitoon et al. 2012; Russell French et al. 2014; Nonhebel and Griffin 2020). This work identified small differences in the expression profiles of *OsYUC9*, *OsYUC11* and *OsYUC12*, suggesting that some subfunctionalisation may occur. *OsYUC12* was found to be exclusively expressed in endosperm for a short period between approximately four and seven days after pollination (DAP). The up- and down-regulation of *OsYUC12* also appeared to coincide with that of *OsIAA29*, encoding an atypical AUX/IAA protein that may play a role as a transcriptional co-regulator in auxin signalling.

A co-expression analysis using *OsYUC12* and *OsIAA29* as bait genes, and online accessed microarray data from several experiments, revealed a small group of genes with the same expression profile restricted to the endosperm from approximately 3 to 7 DAP (Nonhebel and Griffin 2020). This included genes that are exclusively expressed in the dorsal aleurone of rice, *i.e. OsPR602*, *OsPR9a*, *AL1* and *OsNF-YB1* (at the early stage of expression) (Li et al. 2008; Kuwano et al. 2011; Xu et al. 2016). Of these, *OsNF-YB1* is reported to play a key role in endosperm development; plants in which the gene was down-regulated had small grains with chalky endosperm (Xu et al. 2016). The other dorsal aleurone-specific genes, *OsPR602*, *OsPR9a* and *AL1* have been reported to have promoters that direct expression to endosperm transfer cells (Li et al. 2008) or contained *cis*-elements similar to those interacting with ZmMRP-1 (Kuwano et al. 2011), an atypical MYB transcription factor that regulates development of endosperm transfer cells (ETCs) in maize *(Zea mays)* (Gómez et al. 2009). The co-expressed gene group also contained a previously un-reported cluster of MYB-related transcription factor-like genes from rice that are the closest rice homologues to *ZmMRP-1*. These data suggested a possible auxin-signalling network associated with development of cells responsible for uptake of nutrients into the developing endosperm.

Investigations of the role of IAA in rice grain development have been limited by a lack of mutants. More extensive work has been carried out on its importance for the development of maize endosperm, where mutants, *defective endosperm18 (de18)* and *defective kernel18 (dek18)* have reduced expression of IAA biosynthesis genes, reduced IAA content as well as an aberrant basal endosperm transfer layer (BETL) and small, shrivelled grains (Bernardi et al. 2012, 2016). In maize grains, IAA accumulates specifically in the BETL and aleurone (Forestan et al. 2010). Furthermore, treatment of developing maize grains with the auxin transport inhibitor, N-1-naphthylphthalamic acid (NPA), resulted in the formation of a multilayered aleurone instead of a single layer, suggesting that an IAA maximum at the endosperm periphery acts as a signal for aleurone differentiation. The most recent work by Bernardi et al. 2019 investigated the transcriptome of *de18* maize grains. Their results suggest that *ZmMRP-1* as well as genes that are regulated by this transcription factor are down-regulated in the auxin deficient mutant. Thus, IAA may regulate BETL development in maize by controlling expression of *ZmMRP-1.*

We have previously reported evidence for conservation of auxin signalling networks operating during grain development of different cereals (Russell French et al. 2014). Rice does not have a well-defined ETC layer (Hands et al. 2012). However, the dorsal aleurone is likely to play a similar role in apoplastic nutrient transfer to the developing endosperm given its proximity to the vascular trace and enrichment with sugar transporters (Bai et al. 2016; Xu et al. 2016). Based on our previous observations in rice as well as the information from maize, we suggest that IAA may regulate the development of the dorsal aleurone in rice. We therefore hypothesise that expression of *OsYUC12* and *OsIAA29* may be localised to these cells. We tested this hypothesis by investigating, via *in situ* hybridisation, the localisation of *OsYUC12* and *OsIAA29* and comparing this with the previously studied and dorsal aleurone-specific *OsPR602.* To identify precisely the timing of maximum expression of *OsYUC12* and *OsIAA29* as well as test their temporal co-expression with *OsPR602*, *OsPR9a*, *AL1* and three rice homologues of *ZmMRP-1,* here designated as rice *MRP-1-like* or *OsMRPL. (OsMRPL1*, *OsMRPL3* and *OsMRPL4),* we carried out a quantitative expression study from grain samples harvested at daily intervals from 1 to 10 DAP.

The existence of a conserved signalling network within cereals, by which auxin regulates the development of ETCs or cells with ETC-like properties, requires the presence in other cereals of orthologues of key proteins with the same expression profile. As Hands et al. (2012) have previously reported the absence of any orthologues of ZmMRP-1 in other cereals, we investigated the phylogeny and protein structures of ZmMRP-1 and its closest homologues in maize, rice, wheat *(Triticum aestivum)* and *Brachypodium distachyon.* Expression of the wheat *MRP-like* genes was investigated for comparison, using the Wheat Expression Browser (Borrill et al. 2016; Ramírez-González et al. 2018) which has a large number of samples including from dissected grains. A similar phylogenetic and *in silico* expression analysis was carried out on putative cereal orthologues of OsIAA29. Finally, we investigated whether OsIAA29-like proteins are restricted to cereals or whether they occur in dicots and non-grass monocots.

## Materials and Methods

### Plant material and growing conditions

Rice plants (*Oryza sativa* ssp. *japonica* cv. Reiziq) were grown in a greenhouse at the University of New England under natural light with 30° C/18°C day/night temperatures. Seven rice seeds were sown directly into flooded cylindrical plastic pots (50 cm × 15 cm) filled with cracking clay soil (vertosol). When the seedlings reached the 2-3 leaf growth stage, they were thinned to three plants per pot. Plants were watered daily and fertilised fortnightly with the commercial fertiliser Aquasol^®^ (2.0 gm/L) until panicle initiation. Panicles in which approximately half of the spikelets reached anthesis were tagged in the afternoon. The date of tagging was recorded as the day of pollination; the following day was designated as 1 day after pollination (DAP) and so on. Tagged panicles were harvested daily in the afternoon from 1 to 10 DAP. Only superior caryopses were collected, weighed, frozen immediately in liquid nitrogen and stored at −80°C until further use.

### RNA extraction, reverse transcription and quantitative real-time PCR

Total RNA was extracted from 80-100 mg grain samples using Bioline^®^ ISOLATE II RNA Plant Kit (Meridian Bioscience). RNA concentration and purity were measured by NanoDrop™ 8000 Spectrophotometer (Thermo Fisher Scientific). RNA quality was checked by the presence of two clear bands of 18S and 28S rRNAs following agarose gel electrophoresis (Nolan et al. 2006). Only high-quality RNA with A260/A280 ratio in the range of 1.8-2.0 was used in downstream applications.

Transcript sequences of *OsYUC12*, *OsIAA29*, *AL1*, *OsPR602*, *OsPR9a, OsMRPL1*, *OsMRPL3* and *OsMRPL4* were downloaded from PHYTOZOME 12.0 (Goodstein et al. 2012). Primer pairs were designed using Primer3 software (Koressaar et al. 2018) (Supplementary Table S1). Either the left or the right primer of each gene was designed to span an exon-exon boundary in order to avoid amplification of any residual genomic DNA contaminant. Amplification of a single product of the expected size by a primer pair was first confirmed by RT-PCR using Qiagen^^®^^ One-Step RT-PCR kit (Qiagen) followed by agarose gel analysis of the amplified products. The gene for rice ubiquitin-conjugating enzyme E2 (*OsUBC*; *LOC_Os02g42314*) was used as the reference gene (Li et al. 2010).

Quantitative real-time RT-PCR was done in two steps. Bioline^^®^^ SensiFAST^TM^ cDNA Synthesis Kit (Meridian Bioscience) was used to synthesize cDNA from 1.0 μg of total RNA template per reaction according to manufacturer’s instructions. A no-RT control that contained all reaction components except the reverse transcriptase was also included. Bioline^®^ SensiFAST™ SYBR^®^ No-ROX Kit (Meridian Bioscience) was used for the quantitative PCR. Each well of a 96-well plate contained 10 ng of cDNA per 20 μl of final reaction volume. All other reagents were added as per the manufacturer’s instructions. A no-template control was included as negative control. Three biological replicates and two technical replicates were included for each primer set. Reactions were carried out in CFX96 Touch™ Real-Time PCR Detection System (Bio-Rad Laboratories). The amplification program used was as follows: 95°C for 2 min and 40 cycles of 95°C for 5 s, 60°C for 10 s and 72°C for 5 s. Melt curve analysis confirmed the amplification of a single uniform product by each primer pair. This was further confirmed by an agarose gel electrophoresis. Primers with poor amplification efficiency or melt curve analysis implying the presence of more than one product were replaced with new primer sets and the experiment repeated. The manufacturer’s software was used to calculate the expression of the targeted genes relative to the expression of the reference gene. Data from the two technical replicates were first averaged, then the mean and standard error of the three biological replicates from the same developmental stage were calculated.

### *In situ* mRNA hybridisation

*In situ* mRNA hybridisation was carried out using the protocol of Drews 1998 with some modifications. Immature rice grains collected at 5, 6 and 7 DAP were trimmed at both ends, and their palea and lemma were carefully removed. Trimmed grains were immediately fixed in freshly prepared formalin-acetic acid-alcohol (FAA; 3.7% formaldehyde, 5% acetic acid and 50% ethanol) fixative first under gentle vacuum on ice for 15 min and then overnight at 4°C. The fixed grains were dehydrated by a graded ethanol series and xylene before infiltrating them with paraffin wax (Paraplast Plus) in an automated tissue processor (TP1020, Leica Biosystems). The grains were embedded in paraffin wax on an embedding centre (EG1150, Leica Biosystems). Paraffin sections (8.0 μm thick) were cut using a rotary microtome (RM2235, Leica Biosystems) and transferred to glass slides coated with poly-L-lysine (Sigma-Aldrich). The slides were air-dried overnight and stored at 4°C until further use.

Purified cDNAs from *OsYUC12*, *OsIAA29* and *PR602* templates were cloned into pGEM^®^-T vector (Promega) following manufacturer’s instructions (refer to Supplementary Table S2 for primer pairs used to amplify the templates). Gene inserts were amplified from the plasmid by T7 and SP6 primers which annealed to T7 and SP6 promoters flanking the inserts. Purified DNA amplicons with flanking T7 and SP6 promoters were used for *in vitro* synthesis of digoxigenin (DIG)-UTP-labelled single-stranded RNA sense and anti-sense probes by using T7 and SP6 polymerases from a DIG RNA Labelling Kit (Roche). The non-complementary sense probe was used as negative control for each gene. The probes were hydrolysed with 200 mM carbonate buffer at 60°C for 80-90 min to generate 150-200 bp fragments.

Selected paraffin sections were de-waxed with Histoclear (Sigma-Aldrich), and rehydrated with a graded ethanol series and PBS. They were treated with 1.0 μg/mL proteinase K solution for 30 min at 37°C. The sections were dehydrated with a graded ethanol series and probes were applied (320 ng probe in 100 μl hybridisation buffer per slide). Hybridisation was carried out overnight in a humidified box at 55°C. After a series of washes, immunodetection of the DIG-labelled probes was carried out by using an anti-DIG antibody coupled with alkaline phosphatase and a ready-to-use 5-bromo-4-chloro-3-indolyl phosphate (BCIP)/nitro-blue tetrazolium (NBT) solution (Sigma-Aldrich) as the chromogenic substrate for the enzyme. The slides were mounted with an aqueous medium made of glycerol and TE buffer (50% v/v). Images of the sections were acquired by a high-definition slide scanner (NanoZoomer 2.0-RS, Hamamatsu Photonics).

### Phylogenetic analysis and database mining

BLASTP search (Altschul et al. 1997) was conducted using OsIAA29 and ZmMRP-1 peptides as queries against proteomes of selected monocots and eudicots on PHYTOZOME 12.0 (Goodstein et al. 2012) and Ensembl Plants 47 (Kersey et al. 2016). Members of the CCA1-like subgroup of MYB-related transcription factors were selected based on information in Du et al. (2013). Molecular Evolutionary Genetics Analysis (MEGA) version X (Kumar et al. 2018) was used for the phylogenetic analysis. The sequences were aligned by MUSCLE (Edgar, 2004). Phylogenetic trees were constructed by the Maximum Likelihood method based on JTT matrix-based model (Jones et al. 1992). The reliability of each node in the tree was determined by the bootstrap test from 500 replicates (Felsenstein 1985). *Os*IAA29 orthologues were compared with homologues from bamboo and non-grass monocots by multiple sequence alignment using Clustal Omega (https://www.ebi.ac.uk). RNA-seq database of Rice Genome Annotation Project (Kawahara et al. 2013) was used to analyse expression profiles of rice genes. Expression data of *OsIAA29* and *OsMRPL* orthologues in wheat and barley (*Hordeum vulgare*) were retrieved from online RNA-seq databases, Wheat Expression Browser http://www.wheat-expression.com (Borrill et al. 2016; Ramírez-González et al. 2018) and BaRTv1.0 https://bio.tools/BaRTv1.0 (Rapazote-Flores et al. 2019), respectively. Co-expression and *cis*-regulatory element (CRE) analyses were conducted as described previously by Nonhebel and Griffin (2020).

## Results

### Gene expression during the first 10 days after pollination

The expression of *OsIAA29* and *OsYUC12* was compared with that of *OsPR602, OsPR9a, AL1* and three *OsMRPLs (OsMRPL1, OsMRPL3* and *OsMRPL4)* in immature grains from 1 to 10 DAP by qRT-PCR. All genes showed a somewhat similar expression profile (Fig. 1). In particular, all genes except *OsMRPL3* were not expressed until 2 to 3 DAP and expression was maximal at 5 to 6 DAP, before declining to low levels by 9 DAP. *OsMRPL3* followed a similar pattern of up-regulation followed by down-regulation, but this was earlier, with expression detected at 1 DAP and maximal expression at 4 to 5 DAP. *OsMRPL4* had the most restricted expression, with very little transcript detected apart from 5- and 6-DAP samples. On the other hand, *AL1* still continued to be active at 10 DAP. As gene expression was only examined in developing grain samples, global expression of genes in other parts of the plant was investigated using RNA-seq datasets available via the Rice Genome Annotation Project. As shown in Fig. 2, expression was mostly restricted to early grain development with no gene activity found in vegetative tissues, floral tissues, or more mature grains at 25 DAP. However, *OsMRPL3, OsMRPL4* and *OsMRPL5* did show slight expression in anthers.

**Fig. 1.**
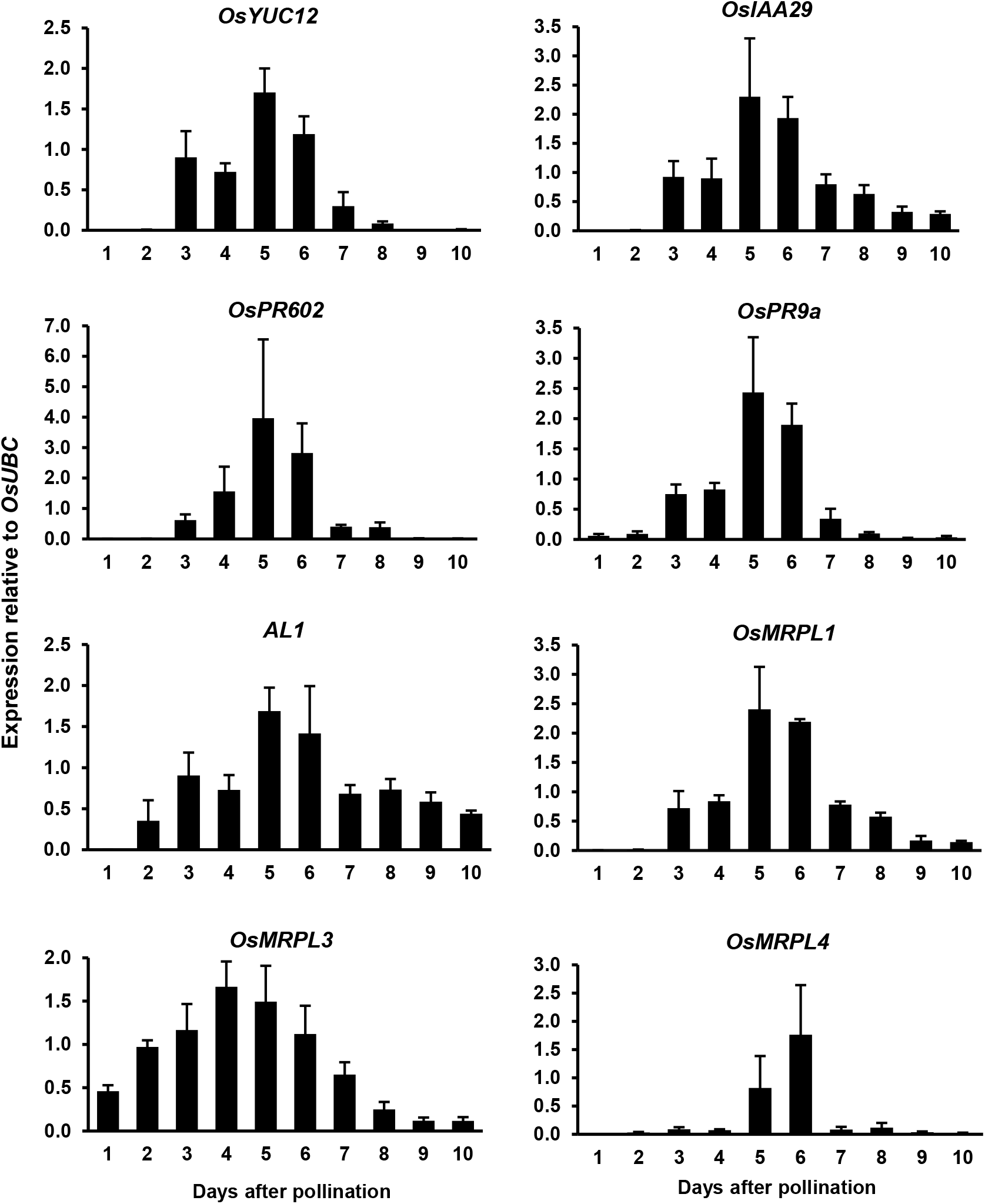
Expression of *OsYUC12, OsIAA29, OsPR602, OsPR9a, AL1, OsMRPL1, OsMRPL3* and *OsMRPL4* in whole rice grains from 1 to 10 days after pollination (DAP). Expression of the genes was calculated relative to the expression level of the reference gene *OsUBC* (*LOC_Os02g42314*), using the software provided by the manufacturer of the CFX96 Touch™ Real-Time PCR Detection System (Bio-Rad Laboratories). Results shown are the means ± the standard errors of the mean (SEMs) of three biological replicates.

**Fig 2.**
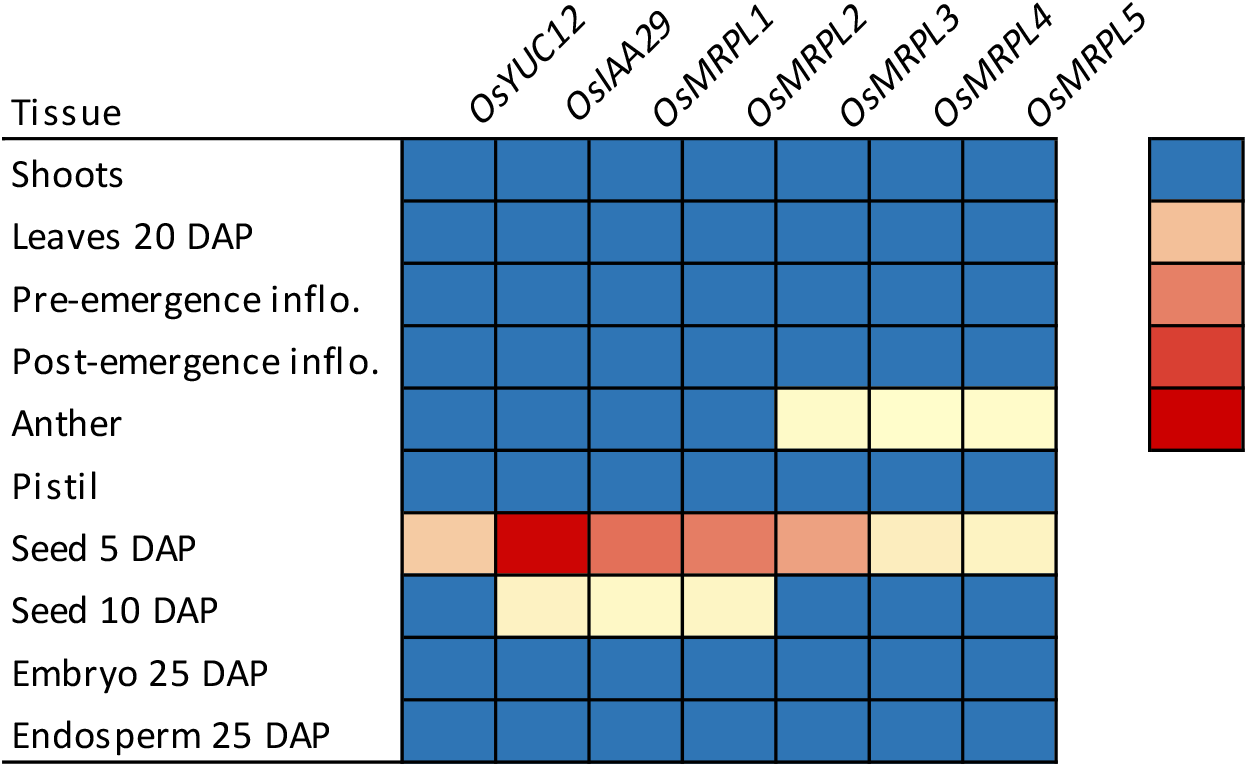
Heat map showing expression of OsYUC12, OsIAA29 and OsMRPL1-5 in different vegetative and reproductive tissues of rice. The expression profiles were retrieved from RNA-seq datasets available on Rice Genome Annotation Project (Kawahara et al. 2013). FPKM = Fragments Per Kilobase of transcript per Million mapped reads.

### Localisation of *OsPR602, OsIAA29* and *OsYUC12* expression in early grains

Spatial expression of *OsPR602, OsIAA29* and *OsYUC12* was examined in immature rice grains by *in situ* mRNA hybridisation. The results reported are from sections of grains collected at 7 DAP, just past the peak of expression, as the larger grains were easier to section and gene expression was still detectable. The sense probes of the genes tested did not give any hybridisation signal, validating the experimentation. Signal from the anti-sense probe of *OsPR602* confirmed its spatial expression in the dorsal aleurone (Fig. 3 b-c and f-g). The spatial expression of *OsIAA29* was similar to that of *OsPR602*, with its hybridisation signal restricted exclusively to the dorsal aleurone (Fig. 4 b-c and f-g). The expression of *OsIAA29* was confined to a narrow strip directly under the major vascular bundle (Fig. 4 f-g). We did not detect any signal for *OsIAA29* expression in the ventral aleurone, starchy endosperm, embryo (Fig. 4d), pericarp and vascular bundles. Spatial expression of *OsYUC12*, on the other hand, followed a much broader pattern than that of *OsPR602* and *OsIAA29*. As shown by both longitudinal and transverse sections, its hybridisation signal was detected in the aleurone and sub-aleurone layers; the signal was distributed both in the dorsal and ventral sides of aleurone (Fig. 5 b-c and f-g). We also detected the hybridisation signal in the embryo (Fig. 5d). However, there was no signal in the starchy endosperm, pericarp and vascular bundles.

**Fig 3.**
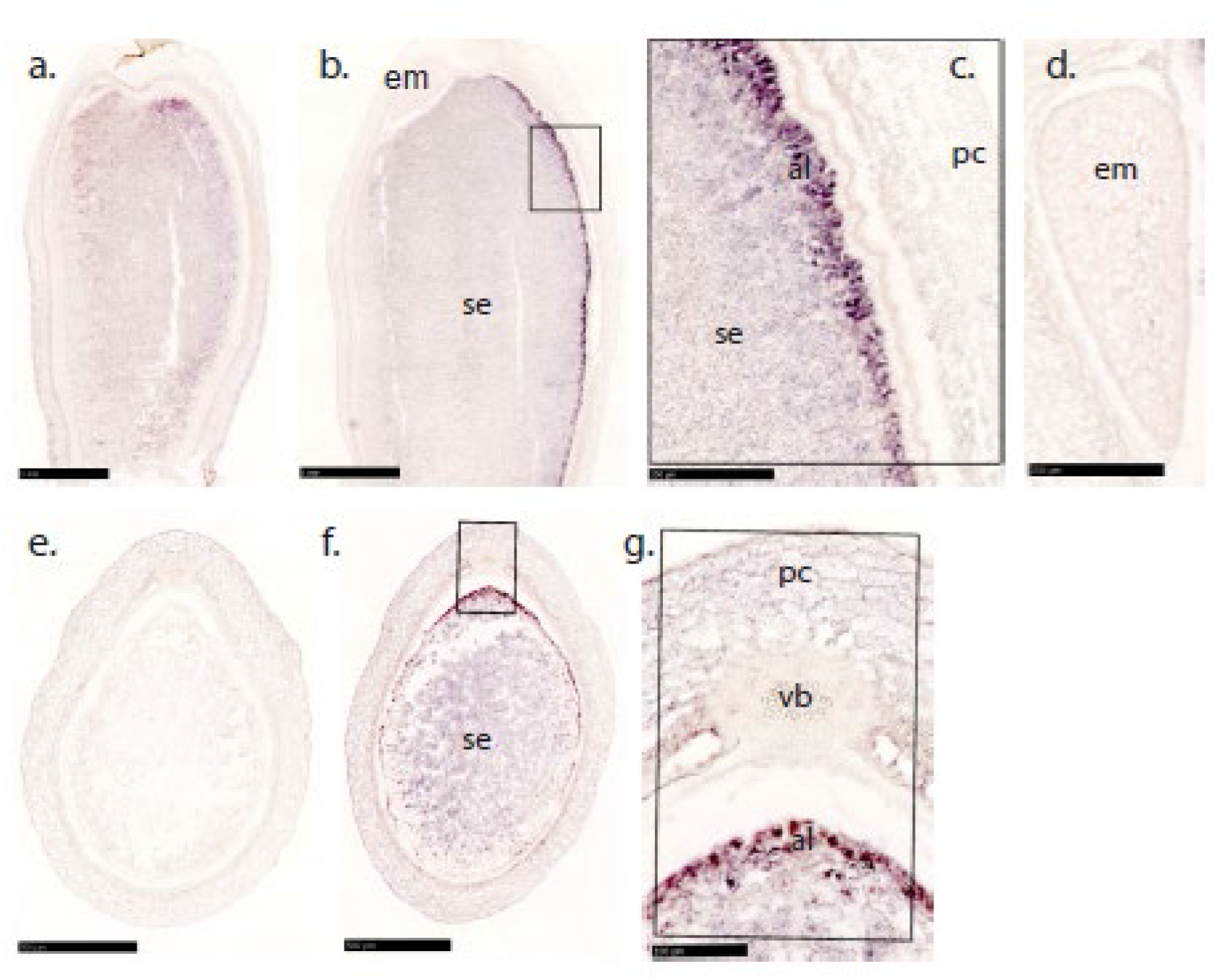
**(a-g).** *In situ* hybridisation of *OsPR602* transcripts in immature rice grains. Hybridisation was done using longitudinal (a-d) and transverse sections (e-g) of immature rice grains at 7 DAP. Images show the absence of *in situ* hybridisation signal from the sense, (a and e) probes and its presence from the anti-sense (b-d and f-g) probes. Image c and g are magnified images of marked areas from b and f, respectively. Image d is derived from the same section as b and shows the absence of hybridisation signal in the embryo. Sense probes (a and e) were used as negative control in the experimentation. Note the strong hybridisation signal in the dorsal aleurone (b-c and f-g). The dorsal side in longitudinal grain sections can be determined by the position of the major vascular bundles running in the pericarp along the dorsal side or by the position of the embryo located always on the ventral side. In case of transverse sections, the dorsal side is determined by the position of the major vascular bundles. Scale bars: (a and b) 1000 μm; (c and d) 250 μm; (e and f) 500 μm; (g) 100 μm. Abbreviations used for annotation: al = aleurone; em = embryo; pc = pericarp; se = starchy endosperm; vb = vascular bundle.

**Fig 4.**
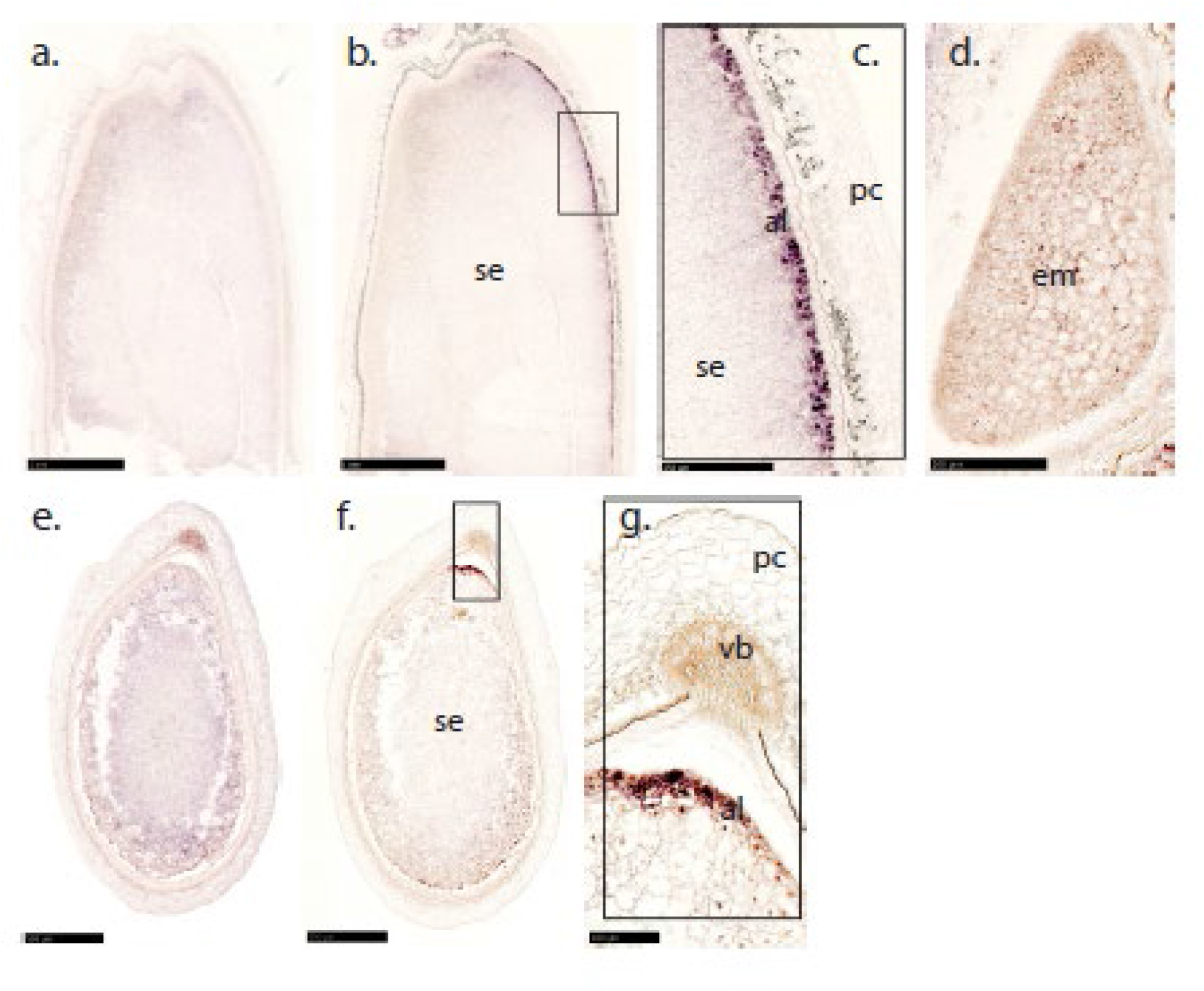
**(a-g).** *In situ* hybridisation of *OsIAA29* transcripts in immature rice grains. Hybridisation was done using longitudinal (a-d) and transverse sections (e-g) of immature rice grains at 7 DAP. Images show the absence of *in situ* hybridisation signal from the sense (a and e) probes and its presence from the anti-sense (b-d and f-g) probes. Image c and g are magnified images of marked areas from b and f, respectively. Image d is derived from the same grain as b and shows the absence of hybridisation signal in the embryo. Sense probes (a and e) were used as negative control in the experimentation. Note the strong hybridisation signal in the dorsal aleurone (b-c and f-g). The dorsal side in longitudinal grain sections can be determined by the position of the major vascular bundles running in the pericarp along the dorsal side or by the position of the embryo located always on the ventral side. In case of transverse sections, the dorsal side is determined by the position of the major vascular bundles. Scale bars: (a and b) 1000 μm; (c) 250 μm; (d) 100 μm; (e and f) 500 μm; (g) 100 μm. Abbreviations used for annotation: al = aleurone; em = embryo; pc = pericarp; se = starchy endosperm; vb = vascular bundle.

**Fig 5.**
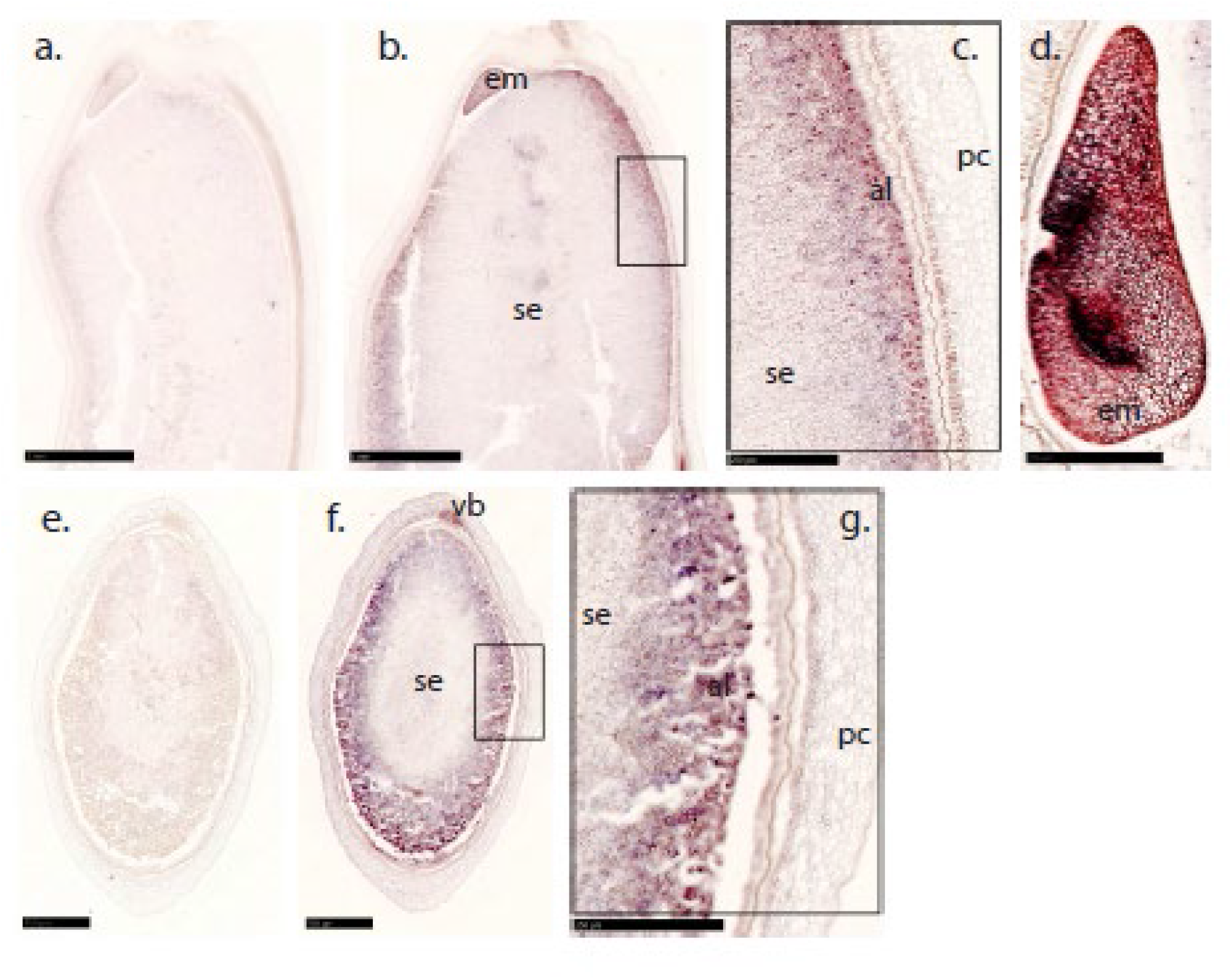
**(a-g).** *In situ* hybridisation of *OsYUC12* transcripts in immature rice grains. Hybridisation was done using longitudinal (a-d) and transverse sections (e-g) of immature rice grains at 7 DAP. Images show the absence of *in situ* hybridisation signal from the sense (a and e) probes and its presence from the anti-sense (b-d and f-g) probes. Image c and g are magnified images of marked areas from b and f, respectively. Image d is derived from the same grain as b and shows the presence of hybridisation signal in the embryo. Sense probes (a and e) were used as negative control in the experimentation. Note the strong hybridisation signal in the aleurone, sub-aleurone and embryo (b-d and f-g). The dorsal side in longitudinal grain sections can be determined by the position of the major vascular bundles running in the pericarp along the dorsal side or by the position of the embryo located always on the ventral side. In case of transverse sections, the dorsal side is determined by the position of the major vascular bundles. Scale bars: (a and b) 1000 μm; (c and d) 250 μm; (e and f) 500 μm; (g) 250 μm. Abbreviations used for annotation: al = aleurone; em = embryo; pc = pericarp; se = starchy endosperm; vb = vascular bundle.

### Phylogenetic analyses of *OsIAA29* and *OsMRPLs* and expression of orthologues in other cereals

As little is known about the non-canonical AUX/IAA protein OsIAA29 or any of its orthologues, we carried out a comprehensive search for, and phylogenetic analysis of similar proteins to explore whether this protein was restricted to cereals. A BLASTP search on PHYTOZOME 12.0 (Goodstein et al. 2012) using OsIAA29 as query against proteomes of selected species found proteins with high homology in both cereal and non-cereal grass species, excepting sequences from moso bamboo *(Phyllostachys edulis* J.Houz.) which showed low peptide homology. Amino acid identity was also low for sequences from non-grass monocots and eudicots. The phylogenetic tree comparing all AUX/IAA proteins from rice with closest homologues from other monocots and eudicots (Fig. 6) confirmed that OsIAA29 had putative orthologues in all grass species tested except moso bamboo. No putative orthologues were found in eudicots. The multiple sequence alignment shown in Suppl. Fig. S1 showed that, like OsIAA29, putative orthologues lacked the N-terminal domains I and II of canonical AUX/IAA proteins and had an acidic C-terminal extension. On the other hand, closest homologous peptides from moso bamboo, banana and the sea grass *Zostera marina* showed typical AUX/IAA domain structure. We explored the expression of barley and wheat orthologues of *OsIAA29* via publicly available RNA-seq databases. The barley orthologue is expressed exclusively in immature grains at 5 DAP, with no expression recorded in embryo at 4 DAP or other tissues (Suppl. Fig. S2A). The expression of wheat orthologues is also restricted to early grain/endosperm around 10 DAP (Suppl. Fig. S2B).

**Fig 6.**
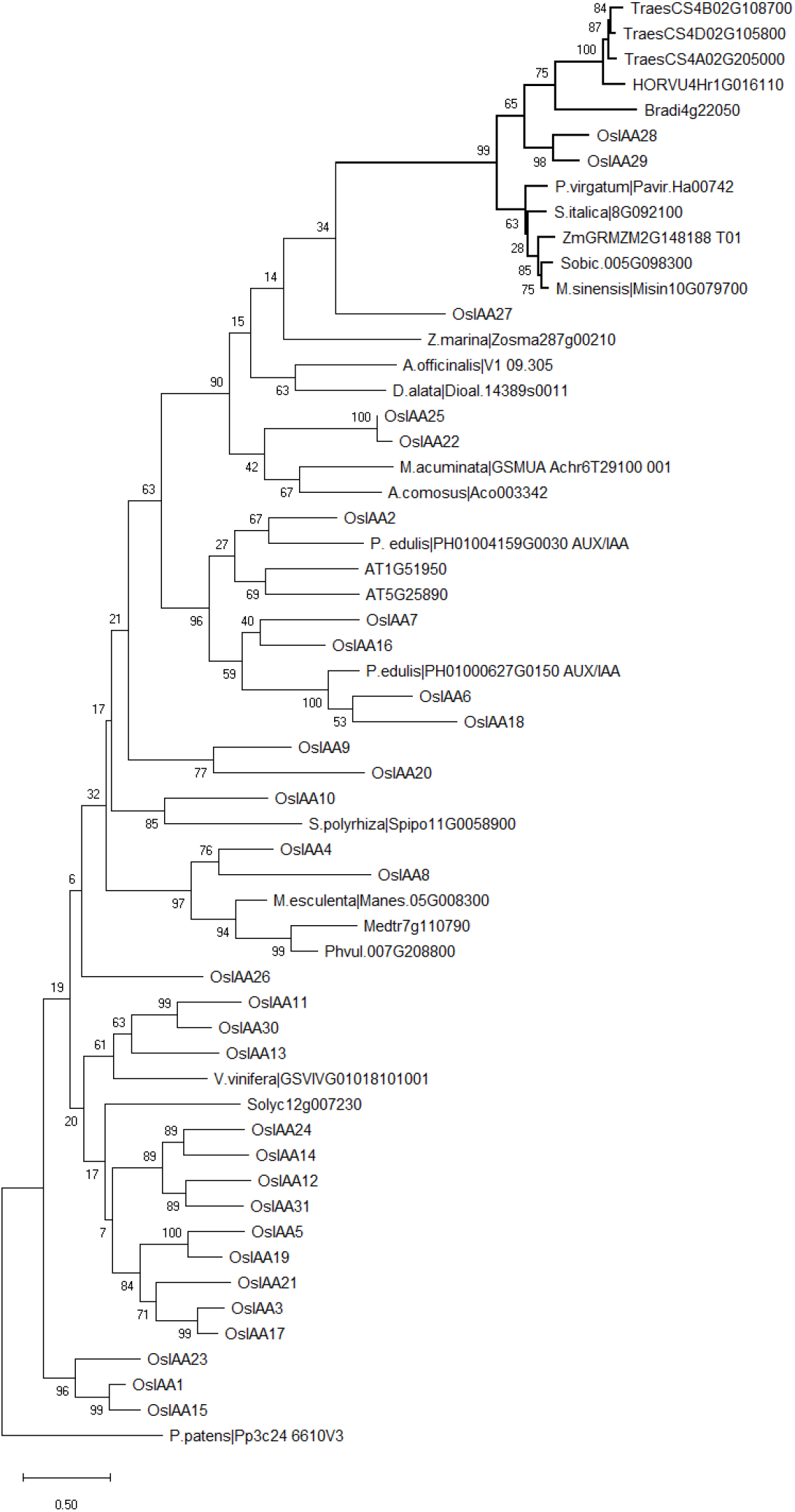
Phylogenetic tree showing relationship of rice AUX/IAA proteins including OsIAA29 with closest homologous peptides from other grasses, non-grass monocots and eudicots. The phylogram was generated by MEGA 10.0 (Kumar et al. 2018), using the Maximum Likelihood method (Jones et al. 1992). Multiple sequence alignments were done using MUSCLE (Edgar 2004). Bootstrap confidence levels were obtained from 500 replicates (Felsenstein, 1985). Monocots included were: *Oryza sativa* (rice), *Phyllostachys edulis* (moso bamboo), *Brachypodium distachyon, Hordeum vulgare* (barley), *Triticum aestivum* (wheat), *Zea mays* (maize), *Sorghum bicolor* (sorghum), *Setaria italica* (millet), *Panicum virgatum* (switchgrass), *Miscanthus sinensis*, *Musa acuminata* (banana), *Ananas comosus* (pineapple), *Dioscorea alata* (purple yam), *Zostera marina* (a sea grass), *Spirodela polyrhiza* (a duckweed) and *Asparagus officinalis* (asparagus). Eudicots included were: *Arabidopsis thaliana*, *Medicago truncatula*, *Phaseolus vulgaris* (common bean), *Solanum lycopersicum* (tomato), *Vitis vinifera* (grape) and *Manihot esculenta* (cassava). The tree is rooted with a sequence from the moss *Physcomitrella patens*. Putative orthologues of OsIAA29 are shown in bold. Scale bar = 0.50 denotes amino acid substitutions per site.

Apart from our previous report of *OsMPRL* genes, no orthologues of ZmMRP-1 have been described. Fig. 7 shows a phylogenetic analysis of all members of the Circadian Clock Associated 1-like (CCA1-like) sub-group of MYB-related transcription factors from maize, rice and *B. distachyon* as well as wheat proteins homologous to ZmMRP-1. The phylogram showed a well-supported clade (bootstrap value of 85) containing ZmMRP-1 and five closely related maize proteins, as well as OsMRPL1-5, four proteins from *B. distachyon* and 21 wheat homologues. As MYB-related transcription factors are highly diverse, the reliability of these relationships was tested by using other methods of phylogenetic analysis: Neighbour Joining (Saitou and Nei 1987) and Minimum Evolution (Rzhetsky and Nei 1992). All analyses reliably grouped the OsMRPLs with ZmMRP-1. All species investigated had multiple paralogues, existing as tandem repeats indicating a high level of recent, lineagespecific gene expansion. OsMRPL1 and OsMRPL2 as well as OsMRPL4 and OsMRPL5 were pairs of tandem duplicates. Although the tree suggests that OsMRPLs as well as proteins from wheat and *B. distachyon* are potentially orthologous to ZmMRP-1, the amino acid identity between OsMRPLs and ZmMRP-1 is very low, ranging from 30 to 38%. However, the multiple sequence alignment (ClustalO) shown in Fig 8 demonstrates homology between ZmMRP-1 and putative orthologues in rice, wheat and *B. distachyon* in the N-terminal domain as well as the MYB domain. It is also noteworthy that ZmMRP-1 itself is missing part of this N-terminal domain compared with other homologues in maize as shown by the inclusion of the sequence encoded by GRMZM2G121111_T01. Although all *OsMRPL* genes appeared to be expressed, a detailed investigation of the peptide sequences (Suppl. Fig. S3) indicated that OsMRPL5 lacked a section of the N-terminal domain found in OsMRPL4 and other proteins. It also had a large insert within the MYB domain. In addition, a comparison of tandem duplicates OsMRPL1 and OsMRPL2 indicated a number of short indels between them.

**Fig 7.**
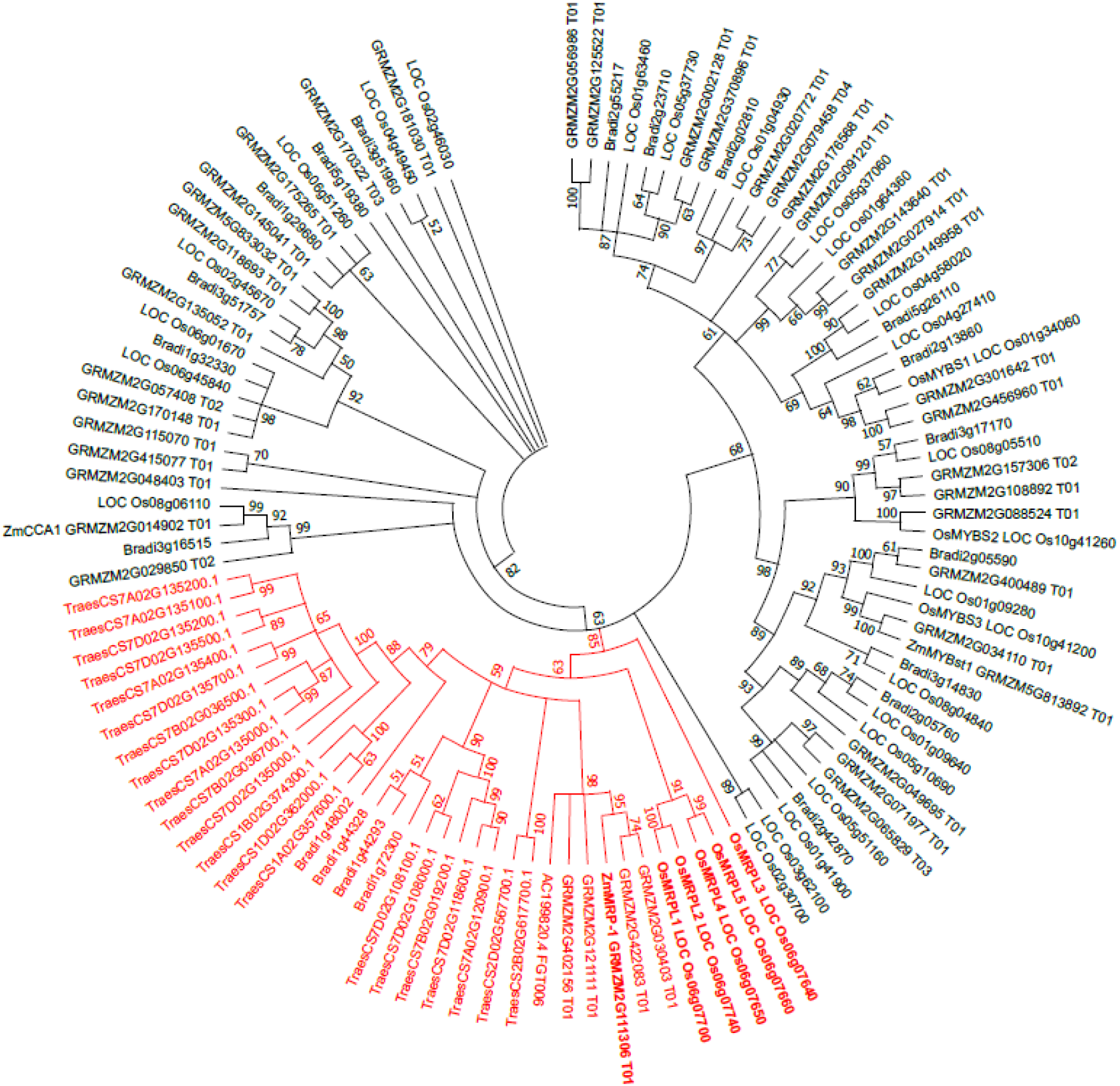
Phylogenetic tree showing the relationships between CCA1-like MYB-related proteins from maize, rice, *Brachypodium distachyon.* and wheat. Due to the large number of wheat proteins, only those in the ZmMRP-1 clade are shown. The consensus phylogram was generated by MEGA 10.0 (Kumar et al. 2018), using the Maximum Likelihood method (Jones et al. 1992). Multiple sequence alignments were done using MUSCLE (Edgar 2004). Bootstrap confidence levels were obtained from 500 replicates (Felsenstein, 1985). The cut-off value for the consensus tree was set at 50. The clade containing ZmMRP-1 is shown in red.

**Fig 8.**
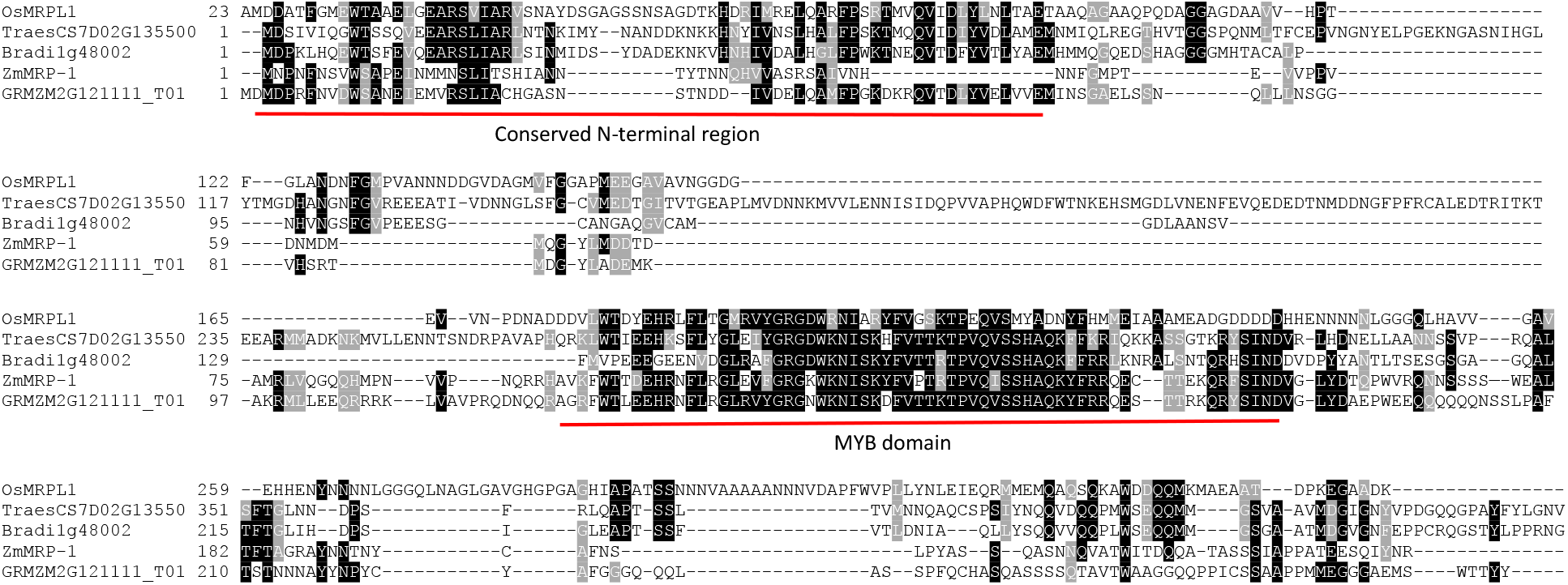
Multiple sequence alignment comparing ZmMRP-1 and one other MRP-1 like protein from maize encoded by GRMZM2G121111_T01 with putative orthologues from rice (OsMRPL1), wheat (TraesCS7D02G135500.1), *B*. *distachyon* (Bradi1g48002). The alignment was carried out in CLUSTAL Omega accessed via https://www.ebi.ac.uk and displayed using Boxshade (https://embnet.vital-it.ch/software/BOX_form.html). Shaded symbols indicate amino acids conserved in at least 50% of sequences.

To investigate possible orthologous expression of MRPL proteins within the grains of another cereal, we used the Wheat Expression Browser as we were unsuccessful in obtaining *in situ* hybridisation results for the *OsMRPLs*. The results shown in Fig. 9 demonstrated expression of 11 out of the 21 wheat *MRPL* genes. These were active exclusively in developing grains with significant expression seen in whole grain samples only at 10 DAP, coinciding with expression of wheat co-orthologues of *OsIAA29*. Although expression in whole grains was very low at 20 DAP, in manually dissected samples, high relative expression was seen exclusively in the endosperm transfer cell (ETC) layer, indicating highly localised up-regulation of these genes. Supplementary Fig. S4 compares expression of *ZmMRP-1* and other full-length members of this clade in maize with that of *ZmIAA38*, the putative orthologue of *OsIAA29*. As with the other cereals there is a strong co-expression seen with maximal expression at 12 DAP.

**Fig 9.**
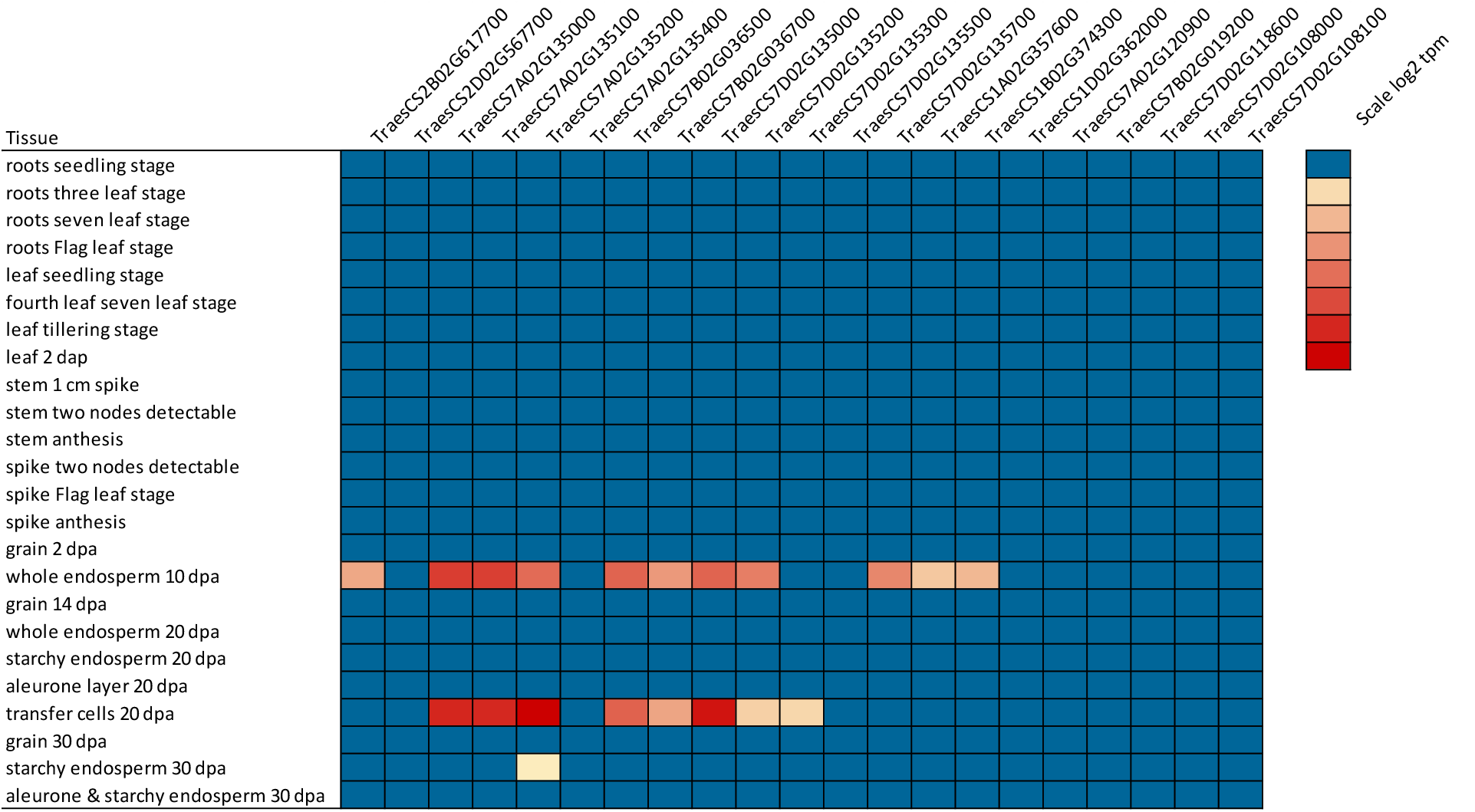
Heat map showing expression of putative wheat orthologues of ZmMRP-1 in different vegetative and reproductive tissues. RNA-seq data is derived from two studies, Wheat development time course (Choulet et al. 2014) and Grain development time course (Pfeifer et al. 2014), both studies using the Chinese Spring variety and accessed via the Wheat Expression Browser (Borrill et al. 2016; Ramírez-González et al. 2018). Gene expression is expressed as log2 TPM (Transcripts Per Million).

### Promoter analysis of genes co-expressed with *OsIAA29, OsPR602* and *OsMRPLs*

The existence of putative orthologues of ZmMRP-1 in rice and other cereals with similar expression profiles suggested conservation of function of these proteins. ZmMRP-1 interacts with a 12-bp *cis*-acting regulatory element (CRE) consisting of two tandem repeats of TATCTC (TC-box) in the promoter region of its target genes (Barrero et al. 2006). With a view to examining the putative enrichment of this CRE in promoter regions of rice genes, we first determined the genes co-expressed with *OsIAA29*, *OsMRPL1* and *OsPR602*, and ranked them as described previously by Nonhebel and Griffin 2020. We then searched for statistically significant enrichment of the CRE in 1000-bp promoter regions of top 300 coexpressed genes, using the RiceFREND platform (Sato et al. 2013). The 12-bp CRE was not found. However, a single TATCTC element was significantly enriched on both promoter strands of the top 100 co-expressed genes (Table 1). Enrichment was strongly associated with co-expression rank.

**Table 1.**
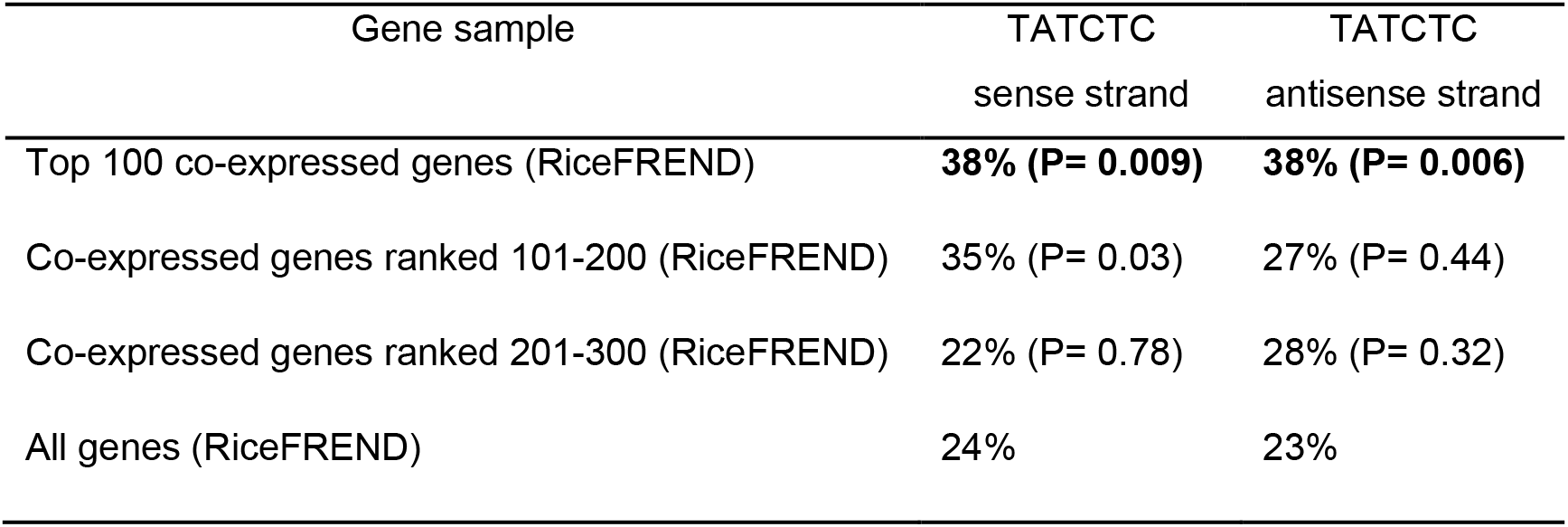
Enrichment of TC-box like TATCTC element in 1000-bp promoter regions of genes coexpressed with *OsIAA29, OsPR602* and *OsMRPL1.* Statistical significance was calculated using Chi-squared tests by comparison to the frequency of this element in all genes (Sato et al. 2013). Its significant enrichment in promoter strands of top 100 co-expressed genes is shown in bold.

## Discussion

In this study, we aimed to investigate IAA biosynthesis and signalling during crucial early stages of endosperm development in rice by exploring in detail the temporal and spatial expression of putative IAA biosynthesis and signalling genes, viz. *OsYUC12* and *OsIAA29*, respectively. Spatial expression was compared with that of *OsPR602,* a gene that has previously been reported as expressed exclusively in the dorsal aleurone of developing rice grains. The precise timing of up-regulation was also compared with that of a group of genes previously suggested (Nonhebel and Griffin 2020) to be co-expressed with *OsIAA29* and *OsYUC12, i.e. OsPR9a, AL1* and three homologues of *ZmMRP-1* that we have designated as *OsMRPLs*.

Our results confirmed that all genes were co-expressed, except for *OsMRPL3* which was up-regulated a little earlier. Expression was grain-specific and short-lived, with genes most active between 3 and 7 DAP. Their up-regulation thus coincides with initiation of coenocyte cellularisation, its completion and differentiation of the early aleurone/starchy endosperm, which take place at 3 DAP, 5 DAP and 7 DAP, respectively (Wu et al. 2016). This supports their possible involvement in an auxin-related signalling pathway during a key stage of transcriptional reprogramming associated with coenocyte cellularisation and differentiation of the early aleurone and starchy endosperm.

Several genes confirmed as temporally co-expressed with *OsIAA29* have localised expression within the dorsal aleurone in rice (Li et al. 2008; Kuwano et al. 2011). Here, we have also demonstrated dorsal aleurone-specific expression of *OsIAA29*. This part of rice aleurone lying adjacent to the major vascular bundles is enriched with sugar transporters and may play an important role in apoplastic nutrient transfer into developing grains (Bai et al. 2016; Xu et al. 2016). Unlike maize, wheat and barley, rice aleurone lacks a well-defined ETC layer (Hands et al. 2012). However, a number of observations point to an ETC-like identity and activity of the dorsal aleurone. Dorsal aleurone-specific *OsPR602* has orthologues in barley (*END1*) and wheat (*TaPR60*), which are also expressed only in ETCs (Doan et al. 1996; Li et al. 2008; Kovalchuk et al. 2009). A *GUS*-reporter gene coupled with *OsPR602* promoter was expressed exclusively in transgenic barley ETCs (Li et al. 2008). Spatio-temporal expression pattern of *OsIAA29* is almost identical to that of *OsPR602.* This suggests a role for the atypical AUX/IAA protein encoded by this gene in signalling during differentiation of dorsal aleurone cells that regulate apoplastic nutrient transfer from maternal pericarp into developing endosperm.

The temporal co-expression of rice homologues of *ZmMRP-1* provides further evidence for a signalling node regulating development of cells with an ETC-like function. Hands et al (2012) have previously reported that no orthologues of ZmMRP-1 could be identified in temperate cereals or in rice. They further observed that expression of the closest homologue in *B*. *distachyon* (Bradi1g72300) could not be detected in developing grain tissue by reverse transcriptase PCR. The very high sequence diversity seen between MYB-related transcription factors makes it very difficult to predict the existence of orthologues of ZmMRP-1. However, our phylogenetic and multiple sequence analyses of homologues of CCA1-like sub-group of MYB-related transcription factors does indicate the existence of a well-supported clade of similar proteins in rice, wheat and *B*. *distachyon.* Our expression analysis via a combination of reverse transcriptase PCR and RNA-seq data also shows that activity of rice and wheat homologues is restricted to a short period during early grain development and furthermore that expression in wheat is highest or possibly restricted to ETCs. The short-lived nature of expression may have led to lack of detection of Bradi1g72300 by Hands et al. (2012). It is also of note that all *MRPLs* appear to have undergone several episodes of gene expansion, with evidence of recent tandem duplication as well as earlier duplication events. Not all genes in each species are expressed; this could also explain the lack of detectable expression of Bradi1g72300. ZmMRP-1 transactivates BETL-specific genes by interacting with a TC-box motif consisting of two 6-bp tandem repeats (TATCTCTATCTC) in their promoter (Barrero et al. 2006). Publications on *AL1* (Kuwano et al. 2011), *OsPR602* and *OsPR9a* (Li et al. 2008) have already suggested that regulation of gene expression within the dorsal aleurone of rice may be regulated in a conserved manner in cereals. Supporting this we found significant enrichment of the TC-box like motif, TATCTC (one of the tandem repeats found in maize) within the promoter regions of genes co-expressed with *OsIAA29, OsPR602* and *OsMRPL1*. Future research focusing on the question of whether OsMRPLs bind to this *cis*-element and regulate gene expression should shed more light on their functions during early endosperm development.

In maize, a study of gene expression in the auxin-deficient *de18* mutant by Bernardi et al. (2019) has provided evidence for the regulation of *ZmMRP-1* and other members of the gene cluster by IAA. The temporal co-expression of *OsMRPL*s with *OsYUC12* and *OsIAA29* suggests that this regulation may be conserved across different cereals. We have previously suggested that three rice *YUCCA* genes predominantly expressed in grains but with differing patterns of temporal expression point to sub-functionalisation and highly localised IAA production at different stages of endosperm development (Basunia and Nonhebel 2019). We hypothesised that the IAA fraction contributed by *OsYUC12* may regulate the molecular events occurring during early stages of endosperm development such as coenocyte cellularisation and early aleurone differentiation. Further support for this view came from its spatial expression pattern. *In situ* hybridisation signal from *OsYUC12* transcripts was detected in the aleurone and sub-aleurone at 7 DAP (Fig. 5), suggesting localised biosynthesis of IAA in early aleurone. In separate work from our laboratory (Kabir et al. 2020), we have also recently reported highest expression of wheat auxin biosynthesis genes *TaYUC9-B1* and *TaYUC9D1* in aleurone and transfer cells although other genes *TaYUC10-A* and *TaYUC10-D* were most highly expression in the starchy endosperm. Forestan et al. 2010 reported strong IAA signal in the aleurone and BETL of maize kernels; *ZmYUC1* expression occurs predominantly in maize aleurone (Zhan et al. 2015; Doll et al. 2017). All results indicate the production of IAA within the aleurone layer of developing cereals grains. Furthermore, *OsYUC12* expression appears to be regulated by two MADS-box transcription factors, MADS78 and MADS79 (Paul et al. 2020). Their expression peaks at 2 DAP and becomes suppressed by 4 DAP. According to the model of Paul et al., they inhibit *OsYUC12* transcription, allowing thereby the proliferation of the coenocyte nuclei. *OsYUC12* transcription is triggered when *MADS78* and *MADS79* expression is suppressed, allowing biosynthesis of IAA which mediates the transition and progression from the coenocytic to the cellularised endosperm. A critical level of endogenous IAA is also required for the cellularisation to proceed in *Arabidopsis* endosperm (Batista et al. 2019).

Although the *in situ* results for *OsYUC12* and *OsIAA29* suggest the production and function of IAA in the dorsal aleurone, the role of OsIAA29 within an auxin signalling pathway is unclear as this protein and its orthologues in other cereals lacks the N-terminal region required for interaction with the TIR1-like auxin receptor. IAA33, a non-canonical AUX/IAA from *Arabidopsis* lacking the degron and EAR domains like *Os*IAA29, negatively regulates auxin signalling in root tip to maintain root distal stem cell identity (Lv et al. 2020). IAA33 competes with the canonical IAA5 for binding to ARF10 and ARF16. *Os*IAA29 may function in a similar way by competing with a canonical *Os*AUX/IAA protein for binding to one or more, as yet unidentified, *Os*ARFs. *OsIAA29* has orthologues only in cereal and non-cereal grass species. Surprisingly, given the close phylogenetic relationship between the subfamilies Oryzoideae and Bambusoideae, no orthologue was detected in moso bamboo. However, over the course of evolution, bamboo species have acquired enormous morphological variation, including different caryopses (Ruiz-Sanchez and Sosa 2015). The ancestral gene for *OsIAA29* was probably lost in moso bamboo. Published reports on the expression and function of *Os*IAA29 orthologues are scarce. The sorghum orthologue is expressed in immature grains at 4 DAP (Nonhebel and Griffin 2020). Analysis of RNA-seq databases indicate early grain-or endosperm-specific expression of the barley, wheat and maize orthologues (Figs. S2 and S4). This gene thus appears to have evolved independently in the grass family (Poaceae) and acquired a novel and grain-specific function during early endosperm development.

In conclusion, we have confirmed the temporal co-expression of *OsYUC12* and *OsIAA29* with genes that have an association with ETC-like cells, *AL1, OsPR602, OsPR9a, OsMRPL1* and *OsMRPL4*. In addition, we have shown that expression of *OsIAA29* is restricted to a narrow strip in the dorsal aleurone very similar to that reported for *OsPR602* and *OsPR9a.* We suggest this gene may act with OsMRPLs to regulate cell differentiation in this region. Further work is required to confirm the localisation of OsMRPLs as well as details of the signalling pathway. Although *OsYUC12* expression appeared to occur over a broader region of the aleurone to *OsIAA29,* our data suggest the production of auxin within the aleurone as has been observed in maize. Future studies focusing on protein-protein interactions of OsIAA29 with OsAUX/IAAs and OsARFs, and targeted inactivation of these genes will provide more information on auxin signalling during early endosperm development.

## Supplementary material

**Suppl. Table S1:**
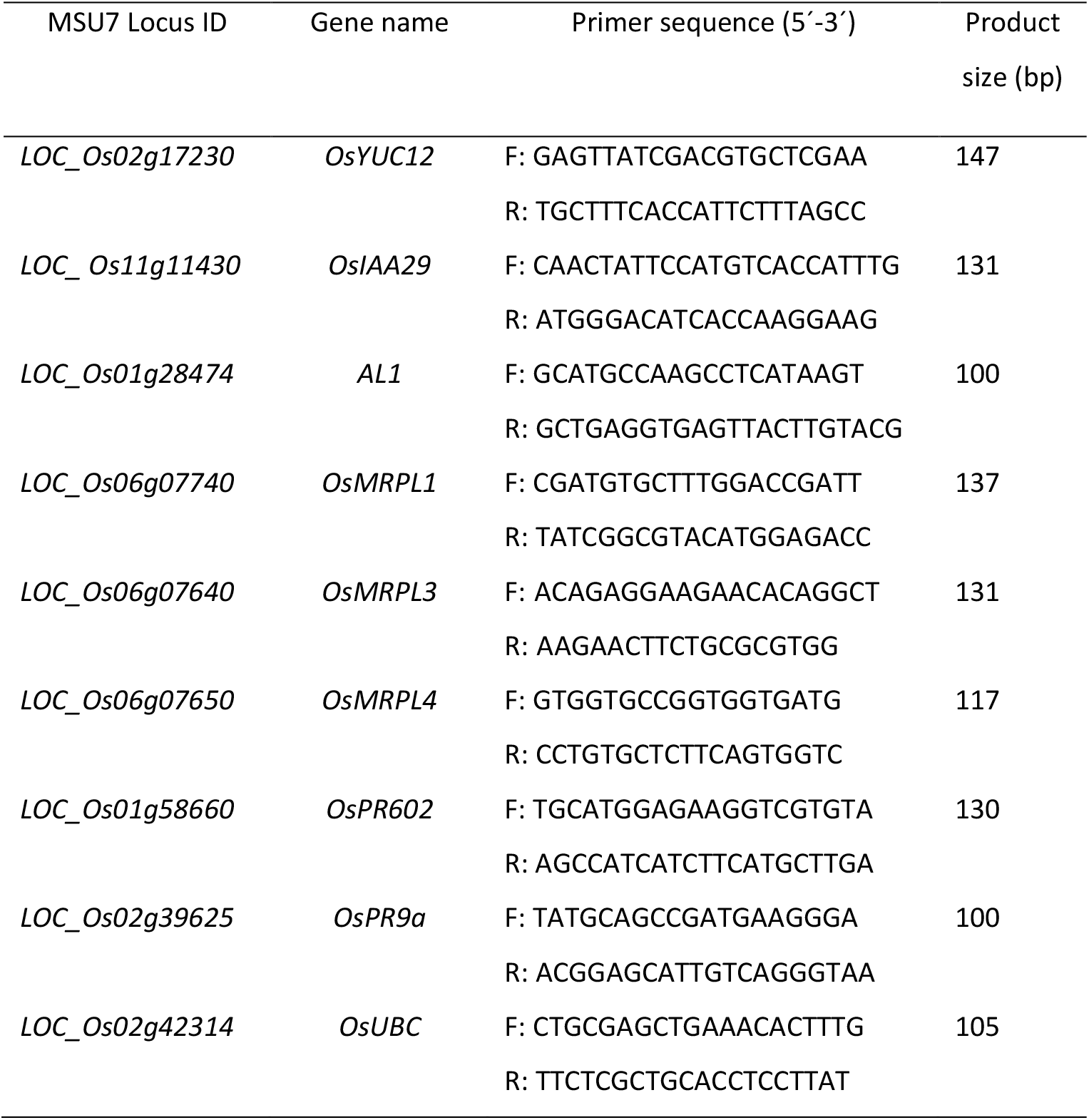
Primer pairs used for qRT-PCR. F=forward primers; R=reverse primers.

**Suppl. Table S2:**
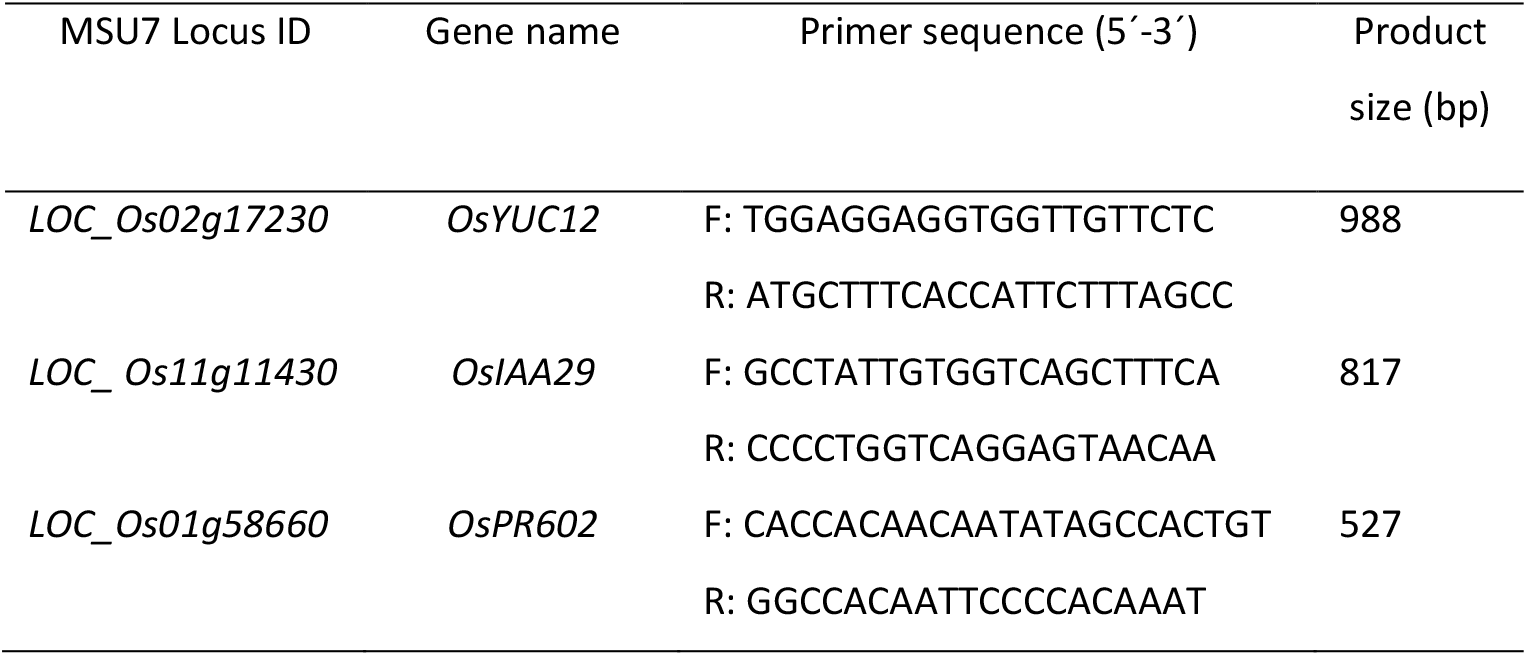
Primer pairs used for probe preparation. F=forward primers; R=reverse primers.

**Suppl. Fig. S1.**
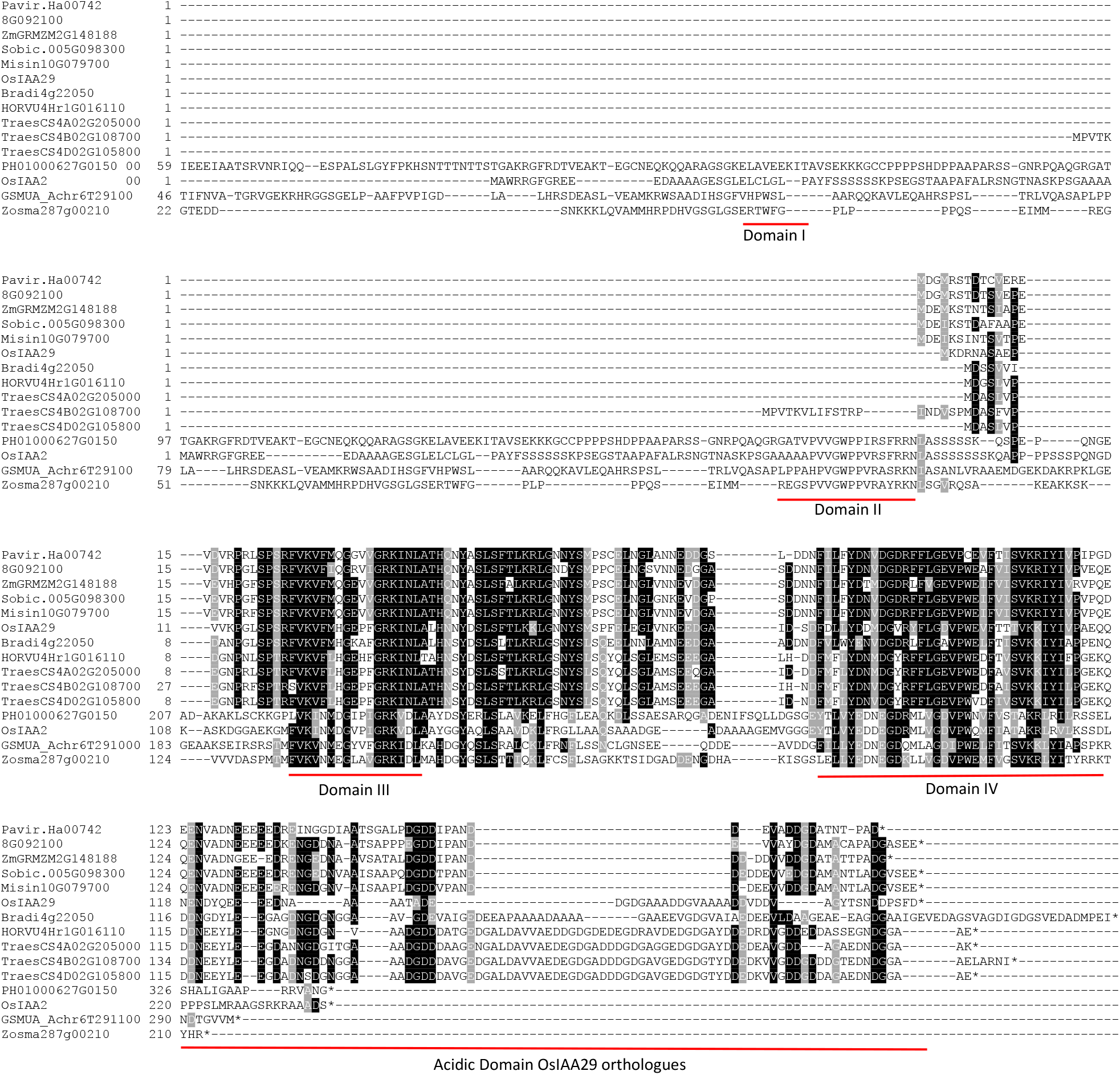
Multiple sequence alignment of OsIAA29, with putative orthologues from other cereals (Pavir.Ha00742 *Panicum virgatum,* 8G092100 *Setaria italica,* ZmGRMZM2G148188_T01 maize, Sobic.005G098300 sorghum, Misin10G079700 *Miscanthus sinensis,* Bradi4g22050 *Brachypodium distachyon,* HORVU4Hr1G016110 barley, TraesCS4B02G108700 etc wheat) and the closest homologous proteins from moso bamboo (PH01000627G0150), banana (GSMUA_Achr6T29100_001) and *Zostera marina* Zosma287g00210.1, as well as canonical AUX/IAA protein OsIAA2. The alignment was carried out in CLUSTAL Omega accessed via https://www.ebi.ac.uk and displayed using Boxshade (https://embnet.vital-it.ch/software/BOX_form.html). Shaded symbols indicate amino acids conserved in at least 50% of sequences.

**Suppl. Fig. S2.**
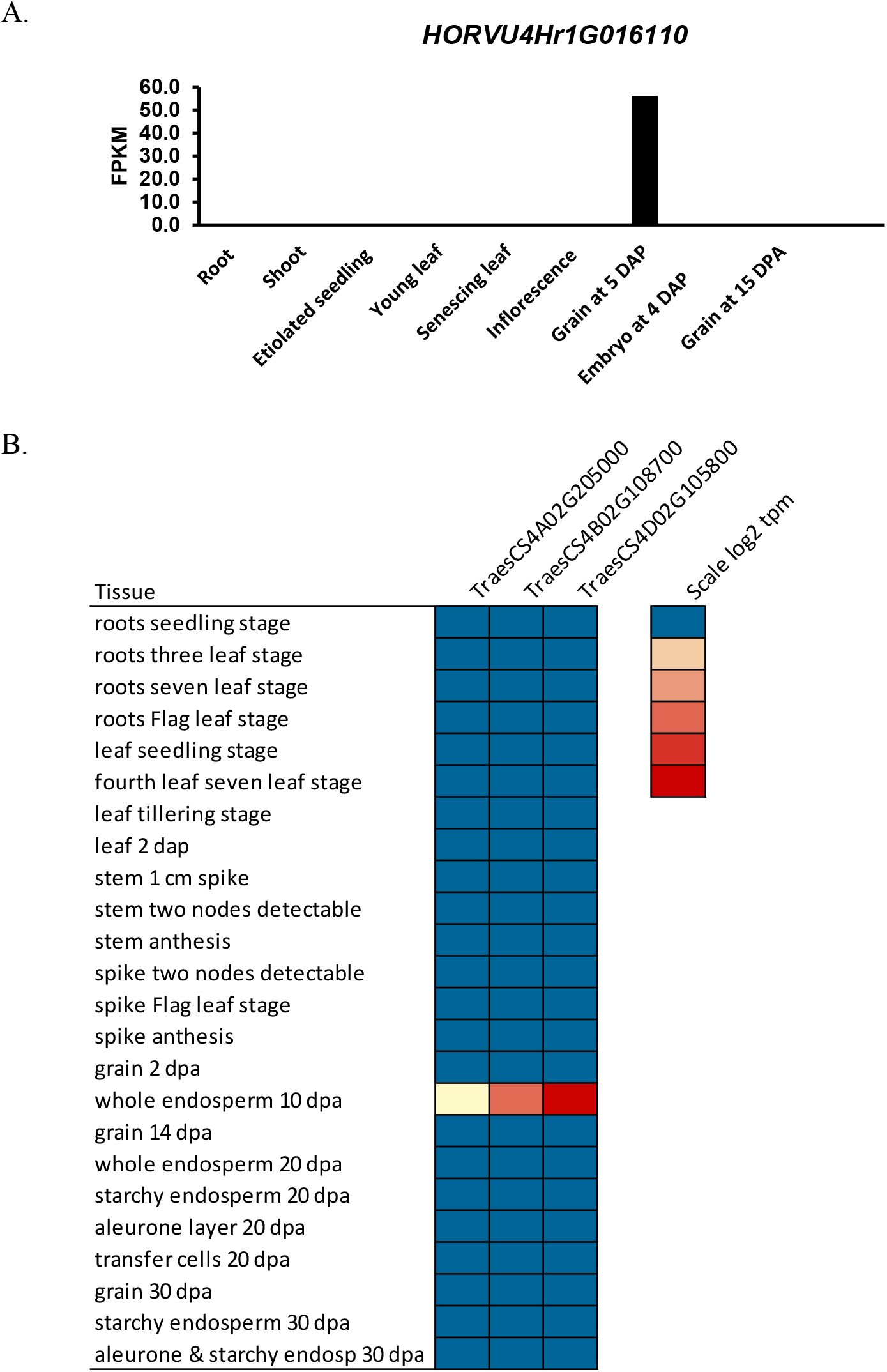
Expression of barley and wheat orthologues of OsIAA29. (A) RNA-seq data for expression of the barley orthologue (HORVU4Hr1G016110) retrieved from BaRTv1.0 (Rapazote-Flores et al. 2019). FPKM = Fragments Per Kilobase of transcript per Million mapped reads. (B) Heat map showing expression of the three wheat orthologues of OsIAA29 in different vegetative and reproductive tissues. RNA-seq data is derived from two studies, Wheat development time course (Choulet et al. 2014) and Grain development time course (Pfeifer et al. 2014), both studies using the Chinese Spring variety and accessed via the Wheat Expression Browser (Borrill et al. 2016; Ramírez-González et al. 2018). Gene expression is expressed as log2 TPM (Transcripts Per Million).

**Suppl. Fig. S3.**
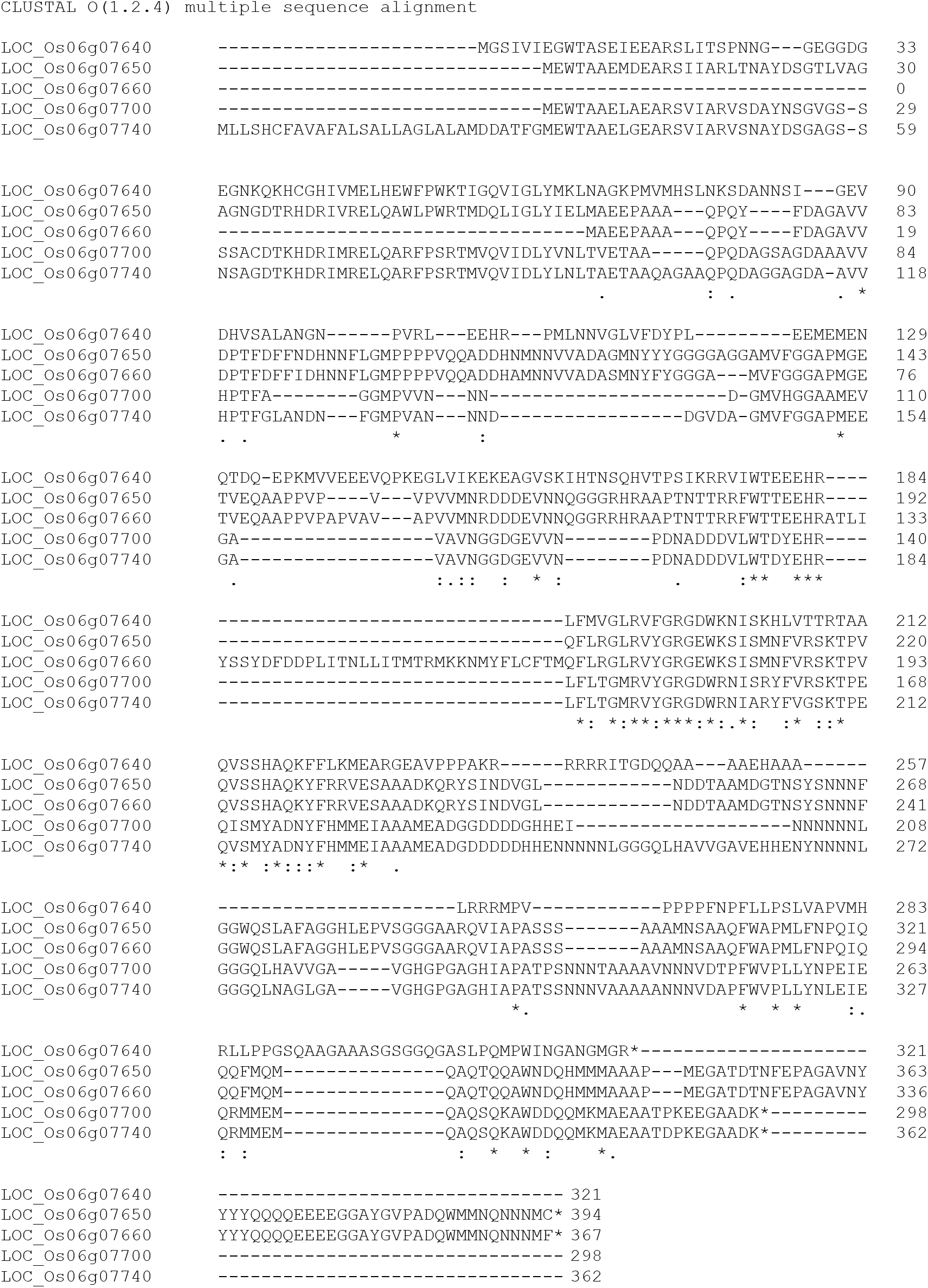
Multiple sequence alignment comparing OsMRPL proteins The alignment was carried out in CLUSTAL Omega accessed via https://www.ebi.ac.uk.

**Suppl. Fig. S4.**
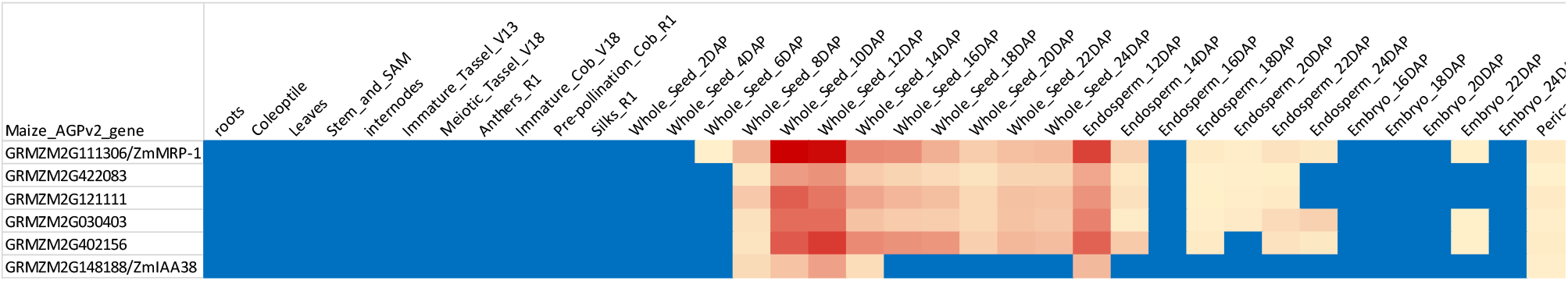
Comparing expression of ZmMRP-1 and related genes with that of ZmIAA38, the maize orthologue of OsIAA29 using RNA-seq data from Stelpflug, et al. (2016). As no expression of any genes was seen in any of the root, leaf, stem or internode samples, a single sample of each only was included in this figure.

